# A putative *erm* gene is present in *ΔermB Clostridioides difficile* isolates showing high levels of resistance to clindamycin

**DOI:** 10.1101/2020.11.20.391128

**Authors:** Viviana Ramírez-Hernández, Gabriel Ramírez-Vargas

**Affiliations:** Hospital “Dr. Rafael Ángel Calderón Guardia”, San José, Costa Rica; Hospital Nacional de Niños “Dr. Carlos Sáenz Herrera”, San José, Costa Rica

**Keywords:** *Clostridiodes difficile*, *erm*, clindamycin

## Abstract

*Clostridioides* difficile has become the leading cause of hospital-acquired diarrhea. Clindamycin is a member of the macrolide-lincosamide-streptogramin B family of protein synthesis inhibitors. Been the ribosomal methylation the most widespread mechanism of resistance to these antibiotics in *C difficile erm* genes usually produce high levels of resistance to these drugs. In this short report we present evidence about the presence of an unreported putative *erm* gene in *C difficile* isolates that despite of presenting a Δ*ermB* genotype maintain high levels of clindamycin resistance.

## Introduction

*Clostridiodes difficile* is a Gram-positive, sporulating and anaerobic bacterium that has become the leading cause of hospital-acquired diarrhea (Banawas, 2018; Guery *et al*, 2019). *C difficile* infection (CDI) is associated with the exposure of the normal intestinal microbiota to antibiotics. The resulting disruption of this normal intestinal microflora allows the multiplication and establishment of *C difficile* in the large intestine causing the pathology (Banawas, 2018; Guery *et al*, 2019).

Antibiotic resistance is crucial in spreading CDI among hospitalized patients, particularly in the older ones (Banawas, 2018). Antibiotics used for treatment of any type of infection could potentially promote (CDI) and resistant to multiple agents may have a selective advantage for the bacteria (Spigaglia, 2016). Most of the antibiotics associated with CDI appearance, including clindamycin, remaining to be associated with the highest risk for CDI (Spigaglia, 2016; Banawas, 2018).

Clindamycin and erythromycin are members of the macrolide-lincosamide-streptogramin B (MLSB) family of protein synthesis inhibitors. Ribosomal methylation is the most widespread mechanism of resistance to these antibiotics in *C difficile. erm* genes usually produce high levels of resistance to these drugs this bymodifying the ribosomal 23S rRNA (Spigaglia, 2016; Banawas, 2018).

The *ermB* gene can be located in the transposon Tn5398, a mobilizable nonconjugative element of 9.6 kb found in *C difficile* genomes and contains two copies of *ermB* (Spigaglia, 2016; Banawas, 2018). Isolates whit a truncated version of Tn5398, containing only one copy of *ermB*, are called Δ*ermB* isolates and fail in inducing the MLS_B_ resistance phenotype (Hussain *et al*, 2005).

In this short report we present evidence about the presence of an unreported putative *erm* gen in *C difficile* isolates that despite presenting a Δ*ermB* genotype maintain high levels of clindamycin resistance.

## Results

**Figure 1.**
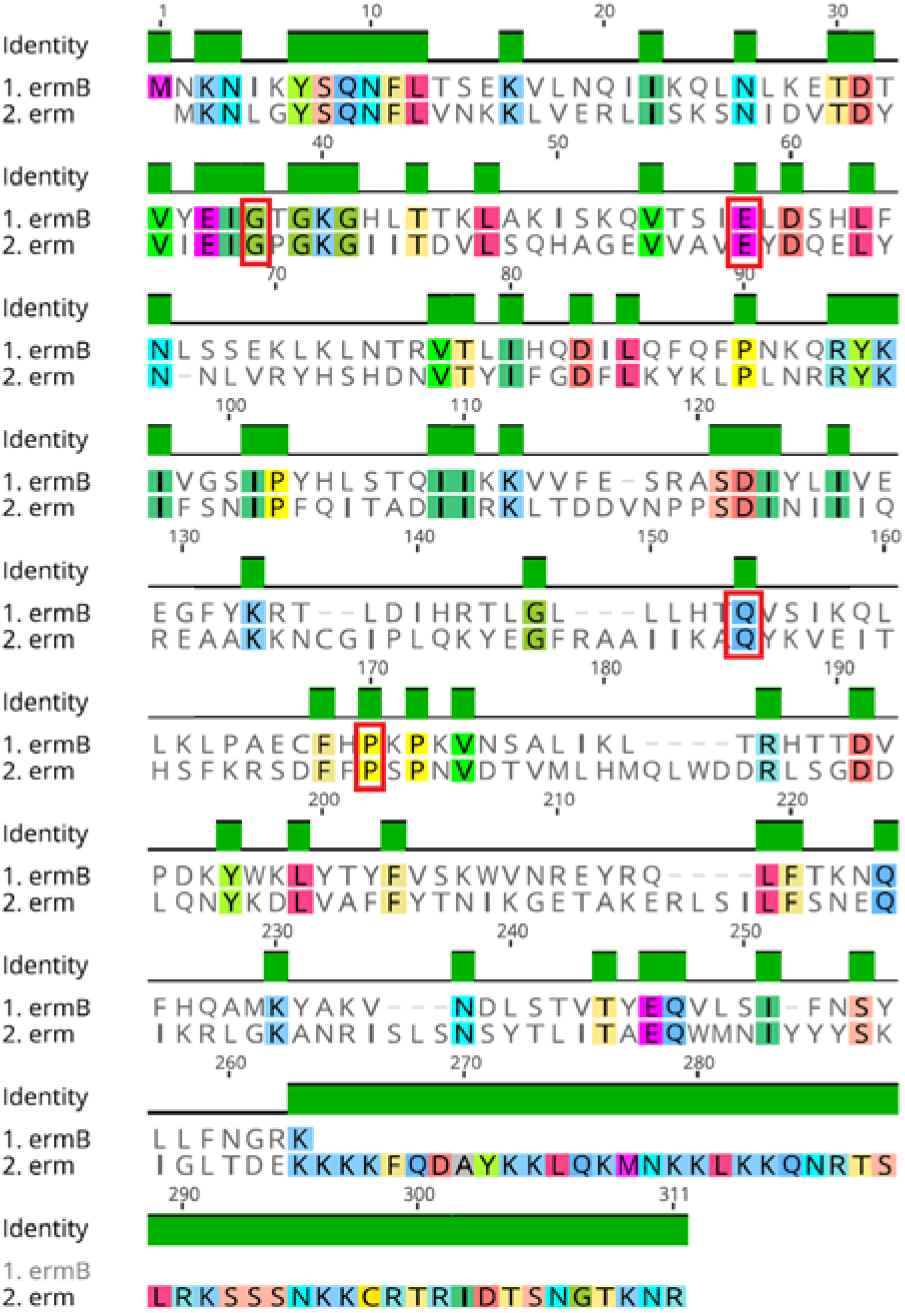
Alignment of *erm* genes. Marked in red boxes are the amino acids associated whit essential ribosome methylation activities.

The gene product analyzed showed a 96% of coverage and 26% identity when compared to ermB from Tn5398. In addition compared to ermB this putative erm protein presents 4 of 6 essential amino acids necessary for ribosome methylation activity already described (Farrow *et al*, 2002).

**Table 1.** Features of analyzed *C difficile* isolates.

| Isolate | *cfrC* | $\Delta ermB$ | CLI MIC<br>($\mu\text{g/ml}$)<br>* |
| --- | --- | --- | --- |
| LIBA-5707 | Yes | Yes | >256 |
| LIBA-5701 | No | Yes | >256 |

\*CLI MIC: clindamycin minimum inhibitory concentration from Ramírez-Vargas *et al*, 2017.

Lacking *cfrC* gene (another ribosome methylating enzyme) and having Δ*ermB* genotype, LIBA-5701is reported to be resistant to clindamycin (MIC>256μg/ml) according to CLSI guides (Ramirez-Vargas *et al*, 2017; Stojković *et al*, 2019).

## Discussion

Here we report the presence of a putative *erm* gen in isolates from Costa Rican Hospitals recovered from CDI patients in the 2000s. The isolates showed similarly pathologic capabilities than other known *C difficile* epidemic isolates like NAP1 (Quesada-Gómez *et al*, 2015; López-Ureña *et al*, 2016).

This putative *erm* gen presents the key amino acids that have been associated to the ribosome methylation activity essential for the high levels of resistance to for MLS_B_ agents such as clindamycinin other erm proteins (Farrow *et al*, 2002). Mutagenesis showed that changes in these amino acids result in lower minimum inhibitory concentration for erythromycin, another agent belonging to the MLS_B_ group (Farrow *et al*, 2002).

In Δ*ermB C difficile* isolates the MLS_B_ resistance phenotype is lost (Hussain *et al*, 2005). So, other resistance mechanism should explain the resistance to clindamycin LIBA-5701. As reported before (Ramirez-Vargas *et al*, 2017; Stojković *et al*, 2019) lack of the element containing *cfrC* did not result in changes in the clindamycin resistance. Using the CARD database (Alcock *et al*, 2020) no other resistance genes associated to MLS_B_ resistance genotype were found (data not shown). At this point we cannot exclude the possibility of other new resistance mechanism snot detected here and the methylation activity of the putative erm gen product must be probe.

## Materials and methods

Raw Illumina SRA files for LIBA-5701 and LIBA-5700used in this work can be found under the accession numbersERR467550andERR467555 respectively at the SRA database. The genomes were assembly using EDENA (Hernandez *et al*, 2008) and annotated using the RAST platform (Aziz, 2008). The predicted erm protein (supplementary material 1) was aligned against ermB from Tn5398 in *C difficle* 630 genome (accession numberNZ_CP019870.1) using Genius platform (Kearse *et al*, 2012). Similarly, they were compared using pBLAST (https://blast.ncbi.nlm.nih.gov/Blast.cgi?PAGE=Proteins) for determination of identity and coverage between the sequences.

Authors declare no conflict of interest.

**Suplemnetary material 1. Putative erm protein sequence in this study** MKNLGYSQNFLVNKKLVERLISKSNID VTDYVIEIGPGKGIITDVLSQHAGEVVA VEYDQELYNNLVRYHSHDNVTYIFGDF LKYKLPLNRRYKIFSNIPFQITADIIRKLT DDVNPPSDINIIIQREAAKKNCGIPLQK YEGFRAAIIKAQYKVEITHSFKRSDFFPS PNVDTVMLHMQLWDDRLSGDDLQN YKDLVAFFYTNIKGETAKERLSILFSNEQ IKRLGKANRISLSNSYTLITAEQWMNIY YYSKIGLTDEKKKKFQDAYKKLQKMNK KLKKQNRTSLRKSSSNKKCRTRIDTSN GTKNR

## References

1. Alcock, B. P., Raphenya, A. R., Lau, T., Tsang, K. K., Bouchard, M., Edalatmand, A., Huynh, W., Nguyen, A. V., Cheng, A. A., Liu, S., Min, S. Y., Miroshnichenko, A., Tran, H. K., Werfalli, R.E., Nasir, J. A., Oloni, M., Speicher, D. J., Florescu, A., Singh, B., Faltyn, M.,… McArthur, A. G. (2020). CARD 2020: antibiotic resistome surveillance with the comprehensive antibiotic resistance database. Nucleic acids research, 48(D1), D517–D525. https://doi.org/10.1093/nar/gkz935

2. Aziz, R. K., Bartels, D., Best, A. A., DeJongh, M., Disz, T., Edwards, R. A., Formsma, K., Gerdes, S., Glass, E. M., Kubal, M., Meyer, F., Olsen, G. J., Olson, R., Osterman, A. L., Overbeek, R. A., McNeil, L. K., Paarmann, D., Paczian, T., Parrello, B., Pusch, G. D.,… Zagnitko, O. (2008). The RAST Server: rapid annotations using subsystems technology. BMC genomics, 9, 75. https://doi.org/10.1186/1471-2164-9-75

3. Banawas S. S. (2018). *Clostridium difficile* Infections: A Global Overview of Drug Sensitivity and Resistance Mechanisms. BioMed research international, 2018, 8414257. https://doi.org/10.1155/2018/8414257

4. Farrow, K. A., Lyras, D., Polekhina, G., Koutsis, K., Parker, M. W., & Rood, J. I. (2002). Identification of essential residues in the Erm(B) rRNA methyltransferase of Clostridium perfringens. Antimicrobial agents and chemotherapy, 46(5), 1253–1261. https://doi.org/10.1128/aac.46.5.1253-1261.2002

5. Guery, B., Galperine, T., & Barbut, F. (2019). *Clostridioides difficile*: diagnosis and treatments. BMJ (Clinical research ed.), 366, l4609. https://doi.org/10.1136/bmj.l4609

6. Hernandez, D., François, P., Farinelli, L., Osterås, M., & Schrenzel, J. (2008). De novo bacterial genome sequencing: millions of very short reads assembled on a desktop computer. Genome research, 18(5), 802–809. https://doi.org/10.1101/gr.072033.107

7. Hussain, H. A., Roberts, A. P., & Mullany, P. (2005). Generation of an erythromycin-sensitive derivative of Clostridium difficile strain 630 (630Deltaerm) and demonstration that the conjugative transposon Tn916DeltaE enters the genome of this strain at multiple sites. Journal of medical microbiology, 54(Pt 2), 137–141. https://doi.org/10.1099/jmm.0.45790-0

8. Kearse, M., Moir, R., Wilson, A., Stones-Havas, S., Cheung, M., Sturrock, S., Buxton, S., Cooper, A., Markowitz, S., Duran, C., Thierer, T., Ashton, B., Meintjes, P., & Drummond, A. (2012). Geneious Basic: an integrated and extendable desktop software platform for the organization and analysis of sequence data. Bioinformatics (Oxford, England), 28(12), 1647–1649. https://doi.org/10.1093/bioinformatics/btsl99

9. Kearse, M., Moir, R., Wilson, A., Stones-Havas, S., Cheung, M., Sturrock, S., Buxton, S., Cooper, A., Markowitz, S., Duran, C., Thierer, T., Ashton, B., Meintjes, P., & Drummond, A. (2012). Geneious Basic: an integrated and extendable desktop software platform for the organization and analysis of sequence data. Bioinformatics (Oxford, England), 28(12), 1647–1649. https://doi.org/10.1093/bioinformatics/bts199

10. López-Ureña, D., Quesada-Gómez, C., Montoya-Ramírez, M., del Mar Gamboa-Coronado, M., Somogyi, T., Rodríguez, C., & Rodríguez-Cavallini, E. (2016). Predominance and high antibiotic resistance of the emerging Clostridium difficile genotypes NAPCR1 and NAP9 in a Costa Rican hospital over a 2-year period without outbreaks. Emerging microbes & infections, 5(5), e42. https://doi.org/10.1038/emi.2016.38

11. Quesada-Gómez, C., López-Ureña, D., Acuña-Amador, L., Villalobos-Zúñiga, M., Du, T., Freire, R., Guzmán-Verri, C., del Mar Gamboa-Coronado, M., Lawley, T. D., Moreno, E., Mulvey, M. R., de Castro Brito, G. A., Rodríguez-Cavallini, E., Rodríguez, C., & Chaves-Olarte, E. (2015). Emergence of an outbreak-associated Clostridium difficile variant with increased virulence. Journal of clinical microbiology, 53(4), 1216–1226. https://doi.org/10.1128/JCM.03058-14

12. Ramírez-Vargas, G., Quesada-Gómez, C., Acuña-Amador, L., López-Ureña, D., Murillo, T., Del Mar Gamboa-Coronado, M., Chaves-Olarte, E., Thomson, N., Rodríguez-Cavallini, E., & Rodríguez, C. (2017). A Clostridium difficile Lineage Endemic to Costa Rican Hospitals Is Multidrug Resistant by Acquisition of Chromosomal Mutations and Novel Mobile Genetic Elements. Antimicrobial agents and chemotherapy, 61(4), e02054–16. https://doi.org/10.1128/AAC.02054-16

13. Spigaglia P. (2016). Recent advances in the understanding of antibiotic resistance in Clostridium difficile infection. Therapeutic advances in infectious disease, 3(1), 23–42. https://doi.org/10.1177/2049936115622891

14. Stojković, V., Ulate, M. F., Hidalgo-Villeda, F., Aguilar, E., Monge-Cascante, C., Pizarro-Guajardo, M., Tsai, K., Tzoc, E., Camorlinga, M., Paredes-Sabja, D., Quesada-Gómez, C., Fujimori, D. G., & Rodríguez, C. (2019). *cfr*(B), *cfr*(C), and a New *cfr*-Like Gene, *cfr*(E), in Clostridium difficile Strains Recovered across Latin America. Antimicrobial agents and chemotherapy, 64(1), e01074–19. https://doi.org/10.1128/AAC.01074-19

